# Gut Microbiome-Driven Metabolites Influence Skin Pigmentation in TYRP1 Mutant Oujiang Color Common Carp

**DOI:** 10.1101/2024.03.01.583018

**Authors:** Roland Nathan Mandal, Jing Ke, Nusrat Hasan Kanika, Xin Hou, Zhiyi Zhang, Penghui Zhang, Huifan Chen, Chunxiao Zeng, Xiaowen Chen, Jun Wang, Chenghui Wang

## Abstract

The gut microbiome is one of the major regulators of the gut-skin axis and is partly regulated by host genetics. In the present study, using comparative high-throughput omics data on CRISPR/Cas9-mediated *TYRP1* mutant and wild-color common carp populations, we quantified the proportion of inter-individual variation in the skin transcriptome and blood metabolome by genetic architecture and gut microbiomes. We found 525 differential metabolites (DMs) and 45 differential gut microbial genera in *TYRP1* mutant fishes relative to the wild type. Through interaction analysis and causal mediation analyses, we revealed that the TYRP1-mutant derived genetic background may exert an inflammatory Acinetobacter - Leukotrience-C4 and – Spermine metabolic pathway under the regulation of an anti-inflammatory cardio-vascular genetic network underlying the upregulating expression of *COMT*, *PLG*, *C2*, *C3*, *F10*, *TDO2*, *MHC1*, and *SERPINF2* gene for evolving unusual coffee-like color phenotype. This unique network appears to underlie the “coffee-like” color phenotype. We propose that the *COMT*-mediated causal effect of the unusual gut microbiome on the atypical skin gene expression patterns through the gut-skin metabolic pathway.

**Article Summary:** R.N. Mandal et al. report on the causal effect of gut microbiome-driven metabolites on the expression pattern of regulatory genes underlying an unusual color phenotype. It suggests that TYRP1 Mutation may rise an unusual inflammatory gut microbiome-skin metabolic pathway that may be balanced by an anti-inflammatory cardio-vascular genetic network leading to unique coloration in Oujiang Color Common Carp.

## Introduction

Teleost fishes have a broader spectrum of skin coloration than birds and mammals because of the combinations of nearly six classes of pigment cells (chromatophores) and variations in the distribution and shapes of these chromatophores (Wang C. et al., 2014; Uwe Irion and Christiane N. V., 2019; Hirata M. et al., 2003). Each of these pigment cells has a specific adaptive role in fish in reflecting the surrounding environmental exposure. For example, located in the basal-most layer melanophores absorb light across the spectrum, iridophores are designed to reflect light, yellow-orange xanthophores in the outermost layer contain pteridine- or carotenoid-based pigments that absorb short-wavelength light, and a continuous sheet of reflective L-iridophores shields the fish body from UV irradiation (Uwe Irion and Christiane N. V., 2019). The most frequent skin color variant relating to eumelanin biosynthesis is in black and brown form (D’Mello, S. A. et al., 2016). In addition to these color variations, different color morphs relating to melanin biosynthesis have been evidenced in fish, birds, mammals, and reptiles, due to natural genetic mutation, such as cinnamon, brown, blond, gray, and white morphs (Uwe Irion and Christiane N. V., 2019; Lewis VM et al., 2019; Jiang, Y., et al., 2014; Puckett, E. E. et al., 2023; Stokowski R. P. et al., 2007).

Skin pigmentation, particularly melanogenesis, can causally be directed by blood metabolites with the help of skin-blood flow through vasoconstriction and vasodilation to the skin under the control of the sympathetic nervous system in the reflection of environmental stress (Garten, J. L., & Fish, F. E., 2020; Plikus, M. V., et al., 2015; Joyner and Dietz, 1997; Yen and Braverman, 1976). Moreover, being heterogeneous in composition blood provides proteins, peptides, signaling molecules, steroid hormones, and other metabolites to various organs, like skin, to facilitate phenotypic characteristics by functioning different metabolic activities of different metabolites (Chen, L. et al., 2022; Bartel J et al., 2015). For example, Astaxanthin was reported to be the cause of yellow and adenine, xanthine and hypoxanthine were of blue color for discus fish (Bo-Tian Yang et al., 2015). A variety of studies extensively analyzed interactions between metabolites in various tissues under different conditions in different species and showed that correlations between circulating blood metabolites are consistent with the known metabolic pathways (Steuer R et al., 2003; Camacho D et al., 2005; Morgenthal K et al., 2005; Krumsiek J et al., 2011).

The condition of an organ and/or tissue, consumed or depleted metabolites from/to blood, may be reflected by the origin and level of specific metabolite shaped by genetic and microbial factors. Several cohort-based studies have already tried to establish in respect of human plasma metabolome that the inter-individual variations can be linked to genetics and the gut microbiome (Asnicar, F. et al., 2021; Bar, N. et al., 2020; Shin, S. Y. et al., 2014; Suhre, K. et al., 2011). It has been reported, for example, in a reference map of potential human serum metabolite determinants, established in 491 individuals from an Israeli cohort, that 182 metabolites are significantly explained by the gut microbiome (Bar, N. et al., 2020). It has also been observed in 1,098 individuals from the United Kingdom and the United States that the diversity of microbial species is associated with favorable cardiometabolic and postprandial markers (Asnicar, F. et al., 2021). Nevertheless, a causal relationship between blood metabolome and gut microbiome exposures directed to skin transcriptome and pigmentation phenotypic outcomes, particularly in fish, is still unresolved.

It is expected that there is a causal relationship between exposure variables, such as blood metabolomes, gut microbiomes, etc., and the skin pigmentation-like phenotypic outcome (Bartel J et al., 2015; Arnold, M. et al., 2015). The Oujiang color common carp (*Cyprinus carpio* var. *color*) under the Cyprinidae family, has been cultured in ponds for over 1200 years in the Oujiang river basin of Zhejiang province in China (Li S., 2003). It has been evident that several variations in skin pigmentation consistently co-exist. However, the classical variety was the fishes with scattered black spots (WB), which could provide an excellent model to explore the underlying molecular mechanisms and their causal inferences of pigment formation and development. Therefore, in our previous study, *TYRP1*, a tyrosinase-related protein 1 encoding gene, was subjected to knocked out to understand its function in melanin biosynthesis (Chen H. et al., 2021). Then, we found that knocking out *TYRP1*gene produced fishes with unusual color variants (Chen H. et al., 2021). In a further mating experiment involving individuals of TYRP1 F0 mutants, we revealed that all the individuals were born with white skin with patches of unusual coffee-like color variant in the F2 generation (**Fig. 1**). In this study, we used high-throughput omics data of skin transcriptome of the skin with coffee-like color phenotype of mutant WB, and the normal black skin of wild fishes of WB variety, blood metabolome, and gut microbiome. We then used multidirectional Causal Mediation analysis to identify the causal inferences of gut microbiome and blood metabolites directed to skin transcriptomic expression underlying unusual coffee-like color phenotype.

**Fig. 1.**
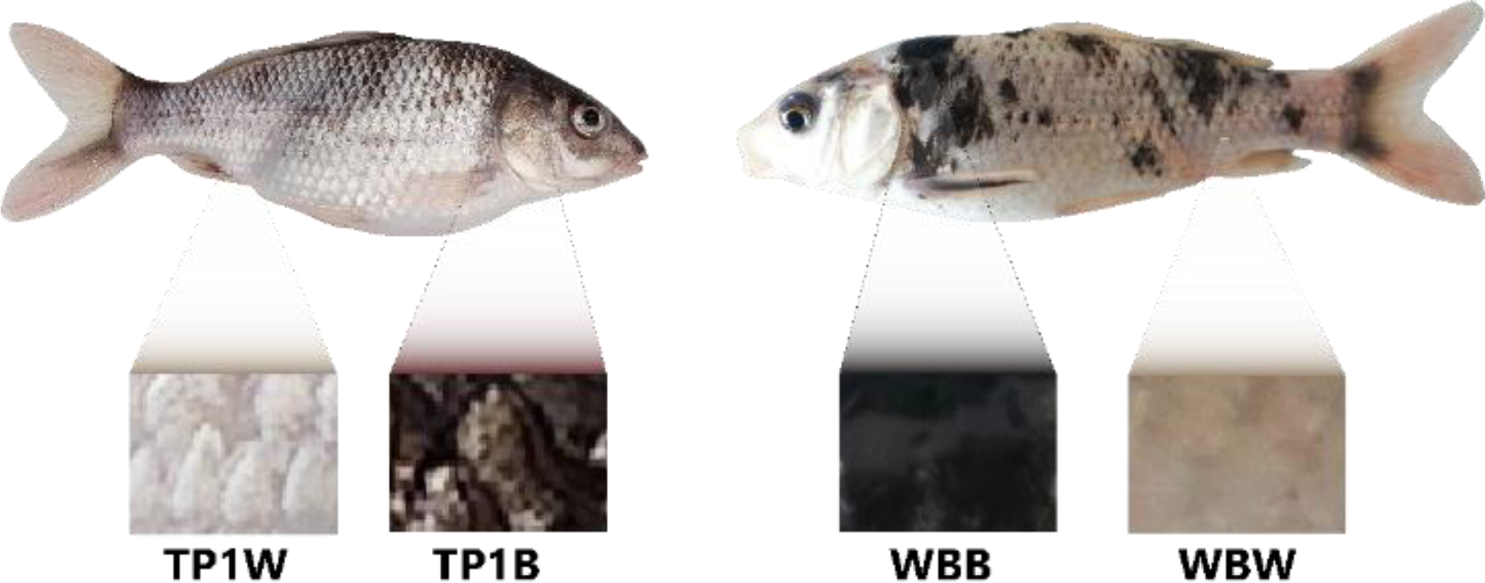
Photograph of TYRP1 Mutant and Wild Type Fishes with Black Patches

## Methodology

### Fish Sampling and Ethical Statement

Both TYRP1 mutant (TYRP1^−/−^), and wild type of white with black variety (WB, TYRP1^+/+^) fishes, reared in a contact pond with controlled water conditions and quality at the Aquatic Animal Genetic Resource Station of Shanghai Ocean University (Shanghai, China), were selected for the study (**Fig. 1**). The *TYRP1* gene was knocked out using CRISPR/Cas9 technology resulting in producing fishes with different color phenotypes, including fishes without black patches from the wild WB. Skin transcriptome data were collected from the skin of four types of color phenotypes (TP1B - skin of coffee-like color of TYRP1 mutant WB, TP1W - white skin of TYRP1 mutant WB, WBB - black skin of wild WB, and WBW - white skin of wild WB) separately from six biological replicates. The sample fishes were temporarily reared and starved in containers for twenty-four hours before dissection. Before dissecting, anesthesia was administered with methane sulfonate (MS222). The dissected skins (2-3cm^2^ in size) were then rapidly poured into liquid nitrogen and stored in a refrigerator at −80°C temperature for RNA extraction. Samples for blood metabolome and gut microbiome were collected from the similar 6 biological replicates. The research was performed as per the ethical guidelines of the Institutional Animal Care and Use Committee (IACUS) of Shanghai Ocean University. We followed the IACUS guidelines to ensure proper care and use of animals during sample collection procedures.

### Differential Metabolomics Analysis

#### Plasma samples analyzed via liquid chromatography–mass spectrometry (LC-MS)

Samples from 6 biological replicates (same individuals used for skin transcriptome) of each group were extracted. After mixing with solvents and internal standards, samples underwent sonication, freezing, and centrifugation to isolate metabolites. Dried samples were then re-dissolved and injected into a Thermo UHPLC-Q Exactive HF-X Mass Spectrometer for LC-MS analysis. Quality control samples were regularly measured throughout the analysis for data reproducibility.

#### Data processing and analysis

Raw data was processed using Progenesis QI software for peak identification, alignment, and filtering. Only metabolites present in over 80% of samples within a group were retained. Missing values were imputed, intensities were normalized, and data was log10 transformed. Normalized data was then compared against HMDB (http://www.hmdb.ca/) and the Metlin (https://metlin.scripps.edu) databases for metabolite identification (Wishart et al., 2018).

#### Statistical analysis

Because of being a high-dimensional and massive feature, we combined univariate and multivariate statistical analysis for screening the differential metabolites between the two biological groups. PCA analysis and PLSDA analysis using ROPLS v1.6.2 software were used to analyze the overall differences between the *TYRP1* Mutant and Wild-Type groups. And fold change and p-value in the univariate analysis, and the volcano map was plotted.

### Gut Microbiome Composition, Diversity, and Discrimination

#### 16S rRNA sequencing revealed gut microbiome composition

DNA from midguts (n = 6 for each group, same individuals used for skin transcriptome and blood metabolome) was extracted by using FastDNA SPIN Kit for Faeces (MP Biomedical) per the manufacturer’s protocols and quantified with a NanoDrop 2000 platform (Shanghai, China). The V3-V4 region of 16S rRNA was amplified and sequenced on an Illumina MiSeq platform. Raw reads were filtered for quality before data processing by using fastp v0.19.6 software (Chen et al., 2018). Paired-end reads were merged and clustered into Operational Taxonomic Units (OTUs) based on 97% sequence similarity (Edgar, 2013). OTU richness and diversity were calculated, and principal coordinate analysis (PCoA) was used to visualize differences between TYRP1 mutant and wild-type groups. Analysis of similarities (ANOSIM) confirmed statistically significant differences in gut microbiome composition (Caporaso et al., 2010).

### Distance matrix-based variance estimation

To understand what drives differences in blood metabolites and gene expression, we performed feature selection. We focused on the 40 most up- and down-regulated metabolites and the 25 hub genes identified in our previous study (unpublished). In this study, we downloaded the RNA-seq data from the CNCB (China National Center for Bioinformation) sequence read archives under the Project accession number PRJCA018085 to calculate the expected TPM for the skin of each color phenotype (WBB, WBW, TP1B, and TP1W). We considered three potential factors: fish phenotype (coffee-like color vs. black patches), genetics (wild-type vs. TYRP1 mutant), and gut microbiome composition. First, we used a PERMANOVA approach to assess the individual contribution of each phenotypic, genetic, and microbial feature to the variation in metabolite levels by applying the adonis function from vegan (version 2.5.5) with 1,000 times permutation (L. Chen et al., 2022). Only features significantly explaining variation (permutation FDR < 0.05) were retained. To address potential redundancy among these features, we used hierarchical clustering to group them based on their correlations. A single representative feature was chosen from each cluster, resulting in a set of non-redundant features. Finally, we used PERMANOVA again to evaluate the combined contribution of these representative features to explain the overall variation in the blood metabolome.

### Microbiome-wide Metabolite Associations

We firstly identified significantly discriminate gut microbiome in both TYRP1 mutant and wild fishes based on their abundance in these two genetic backgrounds. We then discarded the gut microbiome for further interaction analysis because of having the multi-collinearity problem (L. Chen et al., 2022) and selected microbiome having significant power to explain the variability of at least one metabolite concentration. We then applied a general linear regression model for assessing pairwise interactions between the gut microbiome and blood metabolites (L. Chen et al., 2022). Finally, we used Benjamini–Hochberg procedure to adjust the P values.

### Estimating the Variance of Individual Metabolites and Expression of Hub Genes

To estimate the variance of each metabolite and expression of hub genes that were expected to be contributed by genetic and microbial features, we applied machine learning-based lasso regression from the glmnet package (version 2.0.16) (L. Chen et al., 2022). We believe that, while the overall variance explained might be an underestimation of the predictive power of the available data layers, the relative variability explained by each data layer should be representative of the dominant factor that explains most variance in each metabolite. All of the microbial (general species and pathway relative abundance) and genetic features that were significantly associated with a specific metabolite and hub genes at FDR < 0.05 were involved in the model. These features were further selected using lasso with a lambda that gave a minimum mean error from a tenfold cross-validation to control for overfitting and to provide a conservative estimate of model performance. Finally, features selected by lasso were included in the linear model to estimate the variance contributed by different factors, and the adjusted r^2^ and F-test P values were recorded. The FDR was calculated based on the Benjamini–Hochberg procedure.

### Multi-directional Causal Mediation Analysis

For microbial features associated with both metabolites and skin transcriptome (FDR < 0.05), we first checked whether the blood metabolites were associated with the skin transcriptome using Pearson correlation (FDR < 0.05). Next, we carried out a bi-directional mediation analysis with interactions (y = x + m + x × m, where y is the outcome, x is the variable and m represents the mediator) between the mediator and outcome taken into account using the mediate function from mediation (version 4.5.0) to infer the mediation effect of metabolites and transcriptome for microbiome impacts (Tingley D. et al., 2014). The FDR was calculated based on the Benjamini– Hochberg procedure.

### Pathway-based Interaction Network

Through using the MetScape application of Cytoscape_v3.10.0 measured metabolites and genes were mapped onto the corresponding network nodes based on KEGG IDs or HMDB identifiers for metabolites, and Gene IDs for genes. Distances between all mapped pairs of metabolites and transcripts were defined as the shortest path in the network, i.e., the minimal number of reaction steps between them. For instance, a distance of zero between a gene and a metabolite indicates that the metabolite is a potential direct reactant of the reaction catalyzed by the particular enzyme encoded by the gene. A distance of one indicates that the enzyme encoding gene catalyzes a directly connected reaction, which takes a product of the particular metabolite as input, and so on. A distance of infinity (Inf.) was assigned if the respective metabolite and gene were disconnected in the pathway network. Note that the network was treated as undirected, i.e. all reaction directions were ignored.

## Results

### Significantly Co-expressed Hub Genes

In our previous study (unpublished), we applied Weighted Gene Co-expression Network Analysis (WGCNA) and identified twenty-five upregulated and significantly co-expressed genes, term as the hub genes, which were highly correlated (≥ 0.50) with the most significant MEyellow module. According to the KEGG pathway, these genes, including *COMT, PLG, C3, C2, C9, C8B, CFH, CFB, F2, F10, TMPRSS6, APOA1, FGA, FGB, FGG, KNG, TDO2, SERPINC1, SERPIND1, SERPINF2, SERPING1, VTN, ABCB3, C1R*, and *MHC1*, are related to tyrosine metabolism, complement and coagulation cascades, antigen processing and presentation pathways.

### Comparative Differential Metabolites

We identified a total of 525 differential metabolites (DMs) between the *TYRP1* mutant and wild WB (**Fig. 2 and Supplementary Fig. 1**). These metabolites were divided into 6 groups, among which 3 groups were of downregulation and 3 of upregulation in *TYRP1* mutant relative to wild fishes. Among these DMs, Cis-Aconitic acid was significantly upregulated and Protoporphyrin was significantly downregulated anions (**Fig. 2-A-C**). Whereas, Isocitric Acid was significantly upregulated and L-tyrosine was significantly downregulated cations (**Fig. 2-D-F**).

**Fig. 2.**
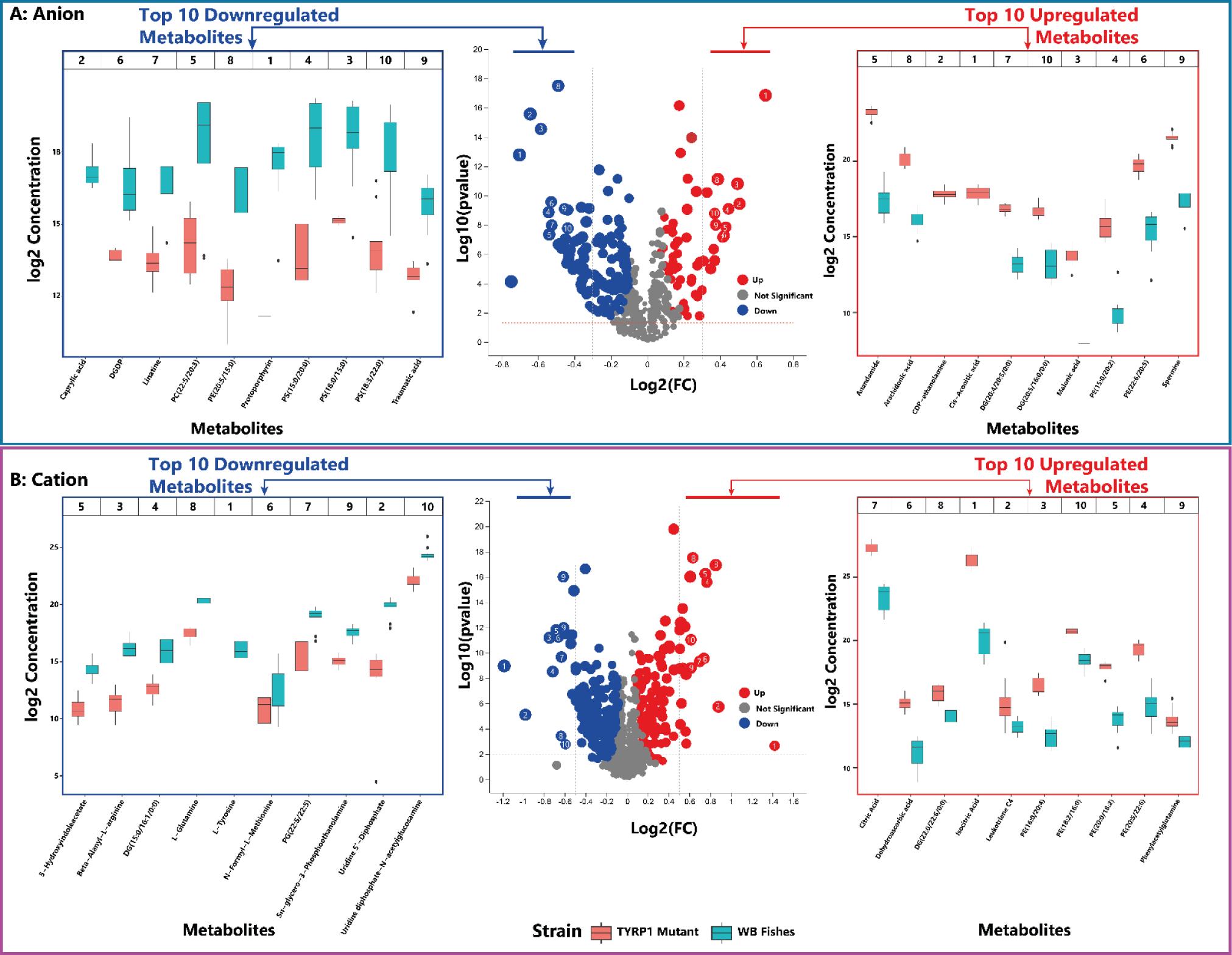
Differentially expressed metabolites in TYRP1 mutant fishes relative to WB fishes. A. Differential metabolites for anions, including top-10 downregulated metabolites (showing number from 1 to 10) at left-side panel and top-10 upregulated metabolites (showing number from 1 to 10) at right-side panel; B. Differential metabolites for cations, including top-10 downregulated metabolites (showing number from 1 to 10) at left-side panel and top-10 upregulated metabolites (showing number from 1 to 10) at right-side panel

### Gut Microbiome Diversity and Discrimination

According to the Simpson’s Index of alpha diversity, several microbial genera were rich in *TYRP1* mutant fishes, thus, microbial populations were more evenly distributed in this assessment group relative to wild fishes (**Supplementary Fig. 2-A**). In the case of beta diversity, the gut microbiomes were far more diversified in *TYRP1* mutant fishes and significantly different (*P_Wilcoxon_ < 0.05*) from wild fishes (**Supplementary Fig. 2-B**). It indicates that the gut microbiome varies between *TYRP1* mutant and wild fish groups. However, 140 microbial genera were identified in both *TYRP1*-mutant and wild fishes. It has been found that 31 and 14 microbial genera were significantly upregulated and downregulated in the *TYRP1* mutant group relative to wild fishes respectively (**Supplementary Fig. 2-C**). The mean distance along with distance ranges on OTU level among *TYRP1* mutant fish individuals was more significant than wild fishes, and even the distance between the two sample groups (**Supplementary Fig. 2-D**). It is, thus, expected that *TYRP1* mutation can influence the abundance and diversity of gut microbiomes. The Wilcoxon Rank test on the abundance of microbial genera at the phylum, family, and genus level showed that Acinetobacter, Micrococcus, Chryseobacterium, Desulfobacterota, Staphylococcus, Rothia, Chloroplast and Competibacter exhibited a significantly higher abundance, and Citrobacter, Neochlamydia, Legionella, Timonella, Kaistia and Galbitalea showed lower abundance in the *TYRP1*-mutant fishes than wild fishes (**Supplementary Fig. 2-E-G**). The distribution pattern of the gut microbial community was examined according to their taxonomy and fishes of two different genetic backgrounds (wild *TYRP1* (^+/+^) and *TYRP1* (^−/−^)) (**Fig. 3**).

**Fig. 3.**
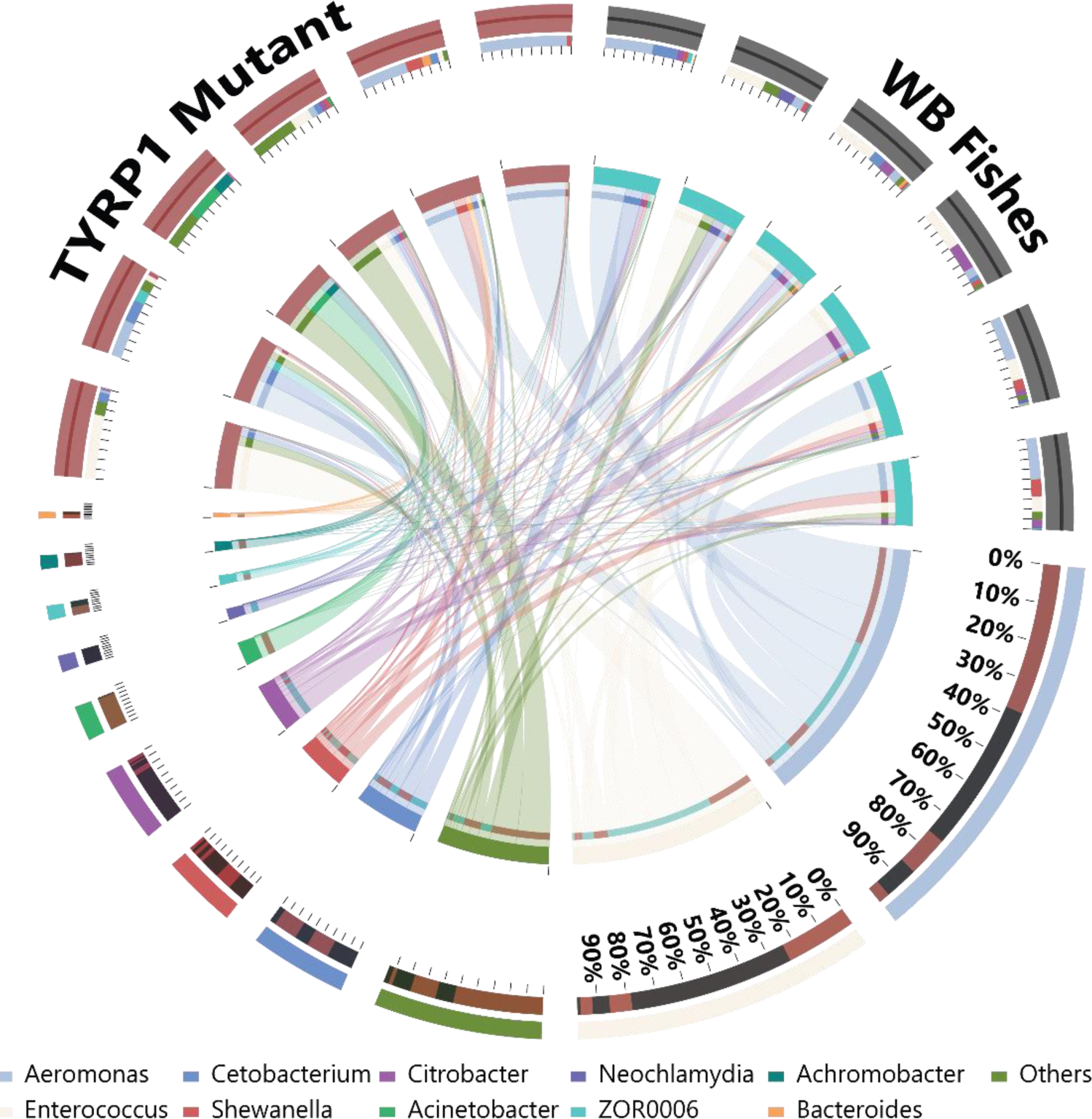
Circos graph for representing the relationship between microbial communities and fishes with two different genetic backgrounds

It has been found that Acinetobacter, Achromobacter, and Bacteroides mainly preferred to be in the *TYRP1* mutant genetic background, whereas Neochlamydia, Citrobacter, and Enterococcus to be in the wild background. Abundance of some microbes, including Aeromonas, and Shewanella, were more or less the same in the two genetic backgrounds. The phylogenetic relationship of the top 50 microbial genera showed that maximum proportions of reads of the *TYRP1* mutant affiliated genera were under Proteobacteria, Firmicutes, and Bacteroidota phyla (**Fig. 4-A**). Among these genera, no credible reads of Acinetobacter and Achromobacter under Proteobacteria, and Chryseobacterium and Pedobacter under Bacteroidota phyla were found in the wild fishes. Furthermore, the linear discriminate analysis (LDA) at the genus level also showed that Acinetobacter, Chryseobacterium, and Macrococcus microbes at the genus level and microbes of Bacteroidota at the phylum level were discriminately abundant in *TYRP1* mutant fishes, whereas Citrobacter and Neochlamydia showed discriminately affinitive to wild fishes at the genus level (**Fig. 4-B**). These results, thus, indicate that the relative abundance of Acinetobacter, Bacteroides, Macrococcus, and Chryseobacterium become highly significant discriminate in *TYRP1* mutant fishes and Citrobacterium, and Neochlamydia in wild fishes.

**Fig. 4.**
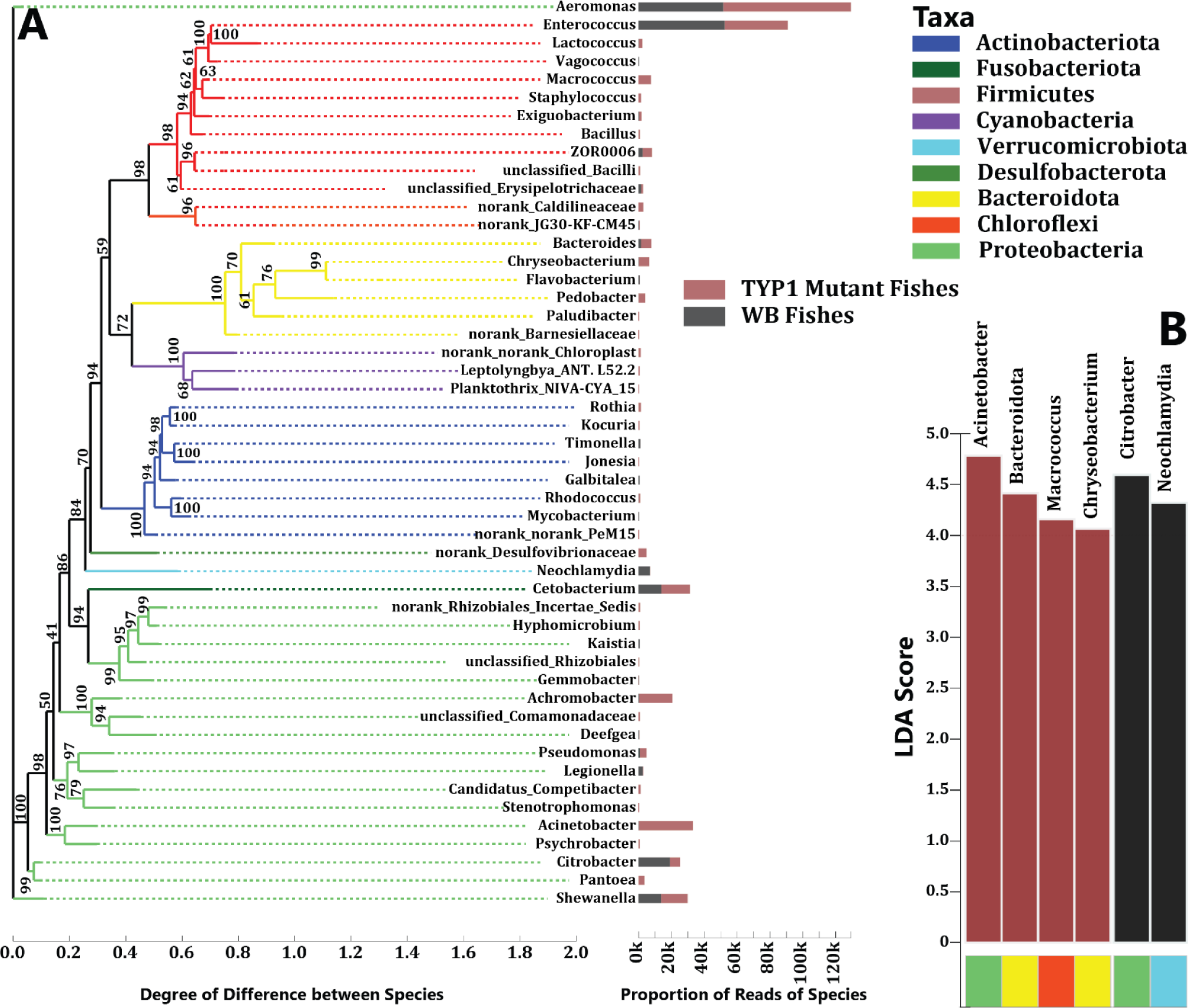
Gut microbiome discrimination in TYRP1 mutant and wild-type fishes. A: Phylogenetic relationship among microbial communities; B: Linear Discriminating Scores of major discriminating microbial genera

#### Tri-active Correlation Network among Skin Transcriptome, Blood Metabolites, and Gut Microbiome

According to the Pearson correlation coefficient, the DMs and DEGs were found to be positively and negatively correlated. The metabolite-metabolites and mRNA-mRNA intra-omics correlation coefficients were comparatively more symmetrically distributed around zero with a maximum absolute correlation coefficient of about 0.53 than inter-omics correlations (metabolite-mRNA) (**Supplementary Fig. 3-A, B**). Moreover, the metabolite-mRNA correlations showed a rather narrower shape, which indicates a tendency to be lower correlations compared to the intra-omics correlations. In contrast, the metabolite-metabolite distribution displayed a wider distribution of correlation coefficients. We then applied Pearson correlation analysis between the top 13 significantly differed gut microbiomes and the top 40 differentially expressed metabolites and found more or less similar distribution pattern compared to intra- and inter-omics correlations with a mentionable difference between metabolite-metabolite and microbiome-metabolite correlation pattern (**Supplementary Fig. 3-C, D**).

The correlation-based network analysis shows that mRNA expression mainly negatively correlated with metabolites and positively with the expression of other mRNA (**Supplementary Fig. 3-E**). However, the *COMT* gene has been found to have a strong positive correlation with Diglyceride (DG) and slightly positive with Arachidonic Acid, with which *TMPRSS6, C8B*, and *CFB* were also positively correlated, but *TDO2, C1R*, and *APOA4* genes were negatively correlated.

We also found strong positive and negative correlations between gut microbiomes and differentially expressed metabolites (**Supplementary Fig. 3-F**). For instance, the highly predominant two microbiomes in *TYRP1* mutant, Acinetobacter, and Achromobacter, have been found to have slightly positive correlations with Spermine, Leukotriene C4, N-Formyl-L-Methionine, Phosphatidylethanolamine and Phenylacetyl-glutamine. L-Tyrosine, the mostly downregulated metabolites in *TYRP1* mutant fishes, was positively correlated with Pseudomonas, Kaistia, Cetobacterium, Enterococcus, Galbitalea, Legionella, Neochlamydia, and Timonella and negatively with Bacteroides, Pedobacter, Citrobacter, Achromobacter, and Acinetobacter.

### Factors Explaining Inter-Omics Variation

Both genetics and gut microbiome variations significantly explained the inter-individual variability of the forty selected metabolomes (top 10 upregulated and downregulated anion and cation metabolites) (**Fig. 5-A)**. It has been found that six gut microbiomes, including Acinetobacter, Citrobacter, Enterococcus, Kaistia, Legionella, and Timonella, had significant power to explain the variability of the differentially expressed metabolites and were significantly associated with a specific metabolite at FDR < 0.05 (**Supplementary Fig. 4**). Among these bacteria, Acinetobacter was significantly associated with 13 blood metabolites, including PG (22:5/22:5) (*R^2^_Adjusted_ = 0.39, P_Pairwise_ = 0.02*), followed by Leukotriene C4 (*R*^2^_Adjusted_ *= 0.38, P_Pairwise_ = 0.02*), X5 Hydroxyindoleacetate (*R^2^_Adjusted_ = 0.37, P_Pairwise_ = 0.02*), L-Tyrosine (*R^2^_Adjusted_ = 0.36, P_Pairwise_ = 0.02*), PS (15:.0/20:0) (*R^2^_Adjusted_ = 0.36, P_Pairwise_ = 0.02*), Spermine (*R^2^_Adjusted_ = 0.35, P_Pairwise_ = 0.02*), Dehydroascorbic acid (*R^2^_Adjusted_ = 0.33, P_Pairwise_ = 0.03*), Isocitric Acid (*R^2^_Adjusted_ = 0.32, P_Pairwise_ = 0.03*), Cis Aconitic acid (*R^2^_Adjusted_ = 0.32, P_Pairwise_ = 0.03*), CDP Ethanolamine (*R^2^_Adjusted_ = 0.32, P_Pairwise_ = 0.03*), Citric Acid (*R^2^_Adjusted_ = 0.32, P_Pairwise_ = 0.03*), Protoporphyrin (*R^2^_Adjusted_ = 0.29, P_Pairwise_ = 0.04*) and PE (20:0/18:2) (*R^2^_Adjusted_ = 0.28, P_Pairwise_ = 0.04*) (**Supplementary Fig. 4 and Fig. 5-B**).

**Fig. 5.**
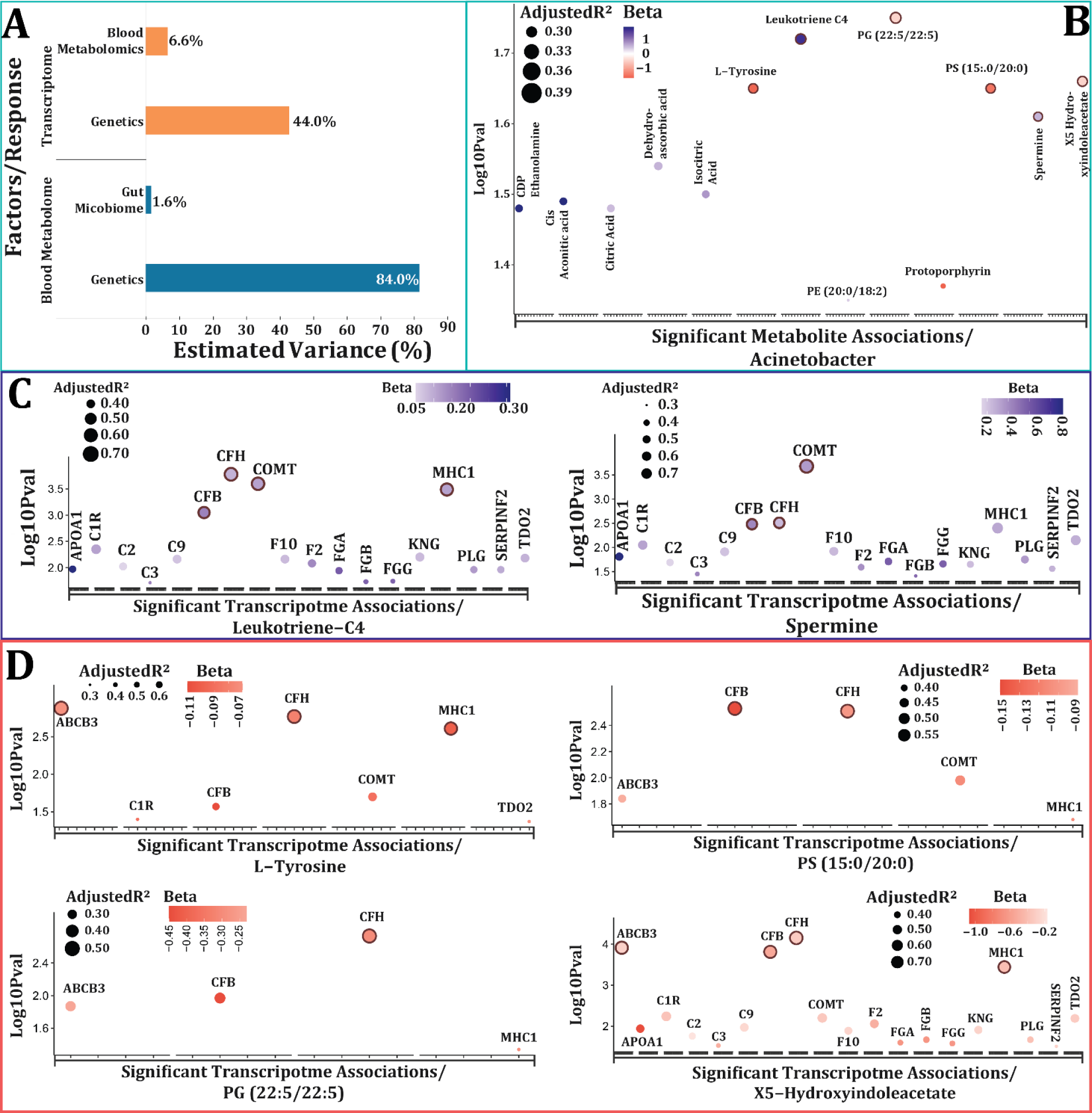
Most significant associations of Acinetobacter-Blood Metabolites-Skin Transcriptomic Expression. A: Factor of explaining the variability of transcriptome and metabolome; B: Most significant associations of Acinetobacter with Blood Metabolites; C: Significant associations of Positive Acinetobacter-associated Blood Metabolite and Skin Transcriptomic Expression; D: Significant associations of Negative Acinetobacter-associated Blood Metabolite and Skin Transcriptomic Expression.

Generalized linear regression model shows that six metabolites among 13 metabolites, including PG (22:5/22:5), Leukotriene C4, X5-Hydroxyindoleacetate, L-Tyrosine, PS (15:.0/20:0), and Spermine were significantly associated with the expression of the 19 hub genes (**Fig. 5-C and – D**). The adjusted R^2^ values indicate that these metabolites can explain 27-79% variability of the hub genes. Among the six metabolites, Leukotriene C4 and Spermine were positively associated with the highest number of hub genes, among which *CFB* (*R^2^_Adjusted_ = 0.65, P_Pairwise_ = 0.001*), *CFH* (*R^2^_Adjusted_ = 0.75, P_Pairwise_ = 0.000*), *COMT* (*R^2^_Adjusted_ = 0.73, P_Pairwise_ = 0.000*), and *MHC1* (*R^2^_Adjusted_ = 0.71, P_Pairwise_ = 0.000*) showed the maximum variability response to Leukotriene C4 and *CFB* (*R^2^_Adjusted_ = 0.55, P_Pairwise_ = 0.003*), *CFH* (*R^2^_Adjusted_ = 0.56, P_Pairwise_ = 0.003*), and *COMT* (*R^2^_Adjusted_ = 0.74, P_Pairwise_ = 0.000*) to Spermine. In contrast, X5-Hydroxyindoleacetate was negatively associated with maximum number of genes and showed maximum power in explaining the variability of *ABCB3* (*R^2^_Adjusted_ = 0.76, P_Pairwise_ = 0.000*), *CFB* (*R^2^_Adjusted_ = 0.75, P_Pairwise_ = 0.000*), and *CFH* (*R^2^_Adjusted_ = 0.79, P_Pairwise_ = 0.000*). Moreover, the interaction analysis found that all six metabolites have the power to explain the expression variability of *MHC1, CFB*, and *CFH* genes. Furthermore, the complement and coagulation pathway-related genes showed positively significant associations with inflammatory Leukotriene-C4 and Spermine metabolites.

### Causal Mediation of Transcriptomic Expression

We finally employed multidirectional causal median analysis to investigate the relationships between 19 significantly associated hub genes, 6 blood metabolites, and Acinetobacter (a bacterium significantly discriminating in *TYRP1* mutant fishes). Both the total effect and average direct effect (ADE) analysis showed a dominant direction: from hub gene expression to blood metabolite concentration mediated by Acinetobacter abundance (e.g., Skin Transcriptome -> Gut Microbiome -> Blood Metabolite). This pattern is evident in **Fig. 6-A and C**.

**Fig. 6.**
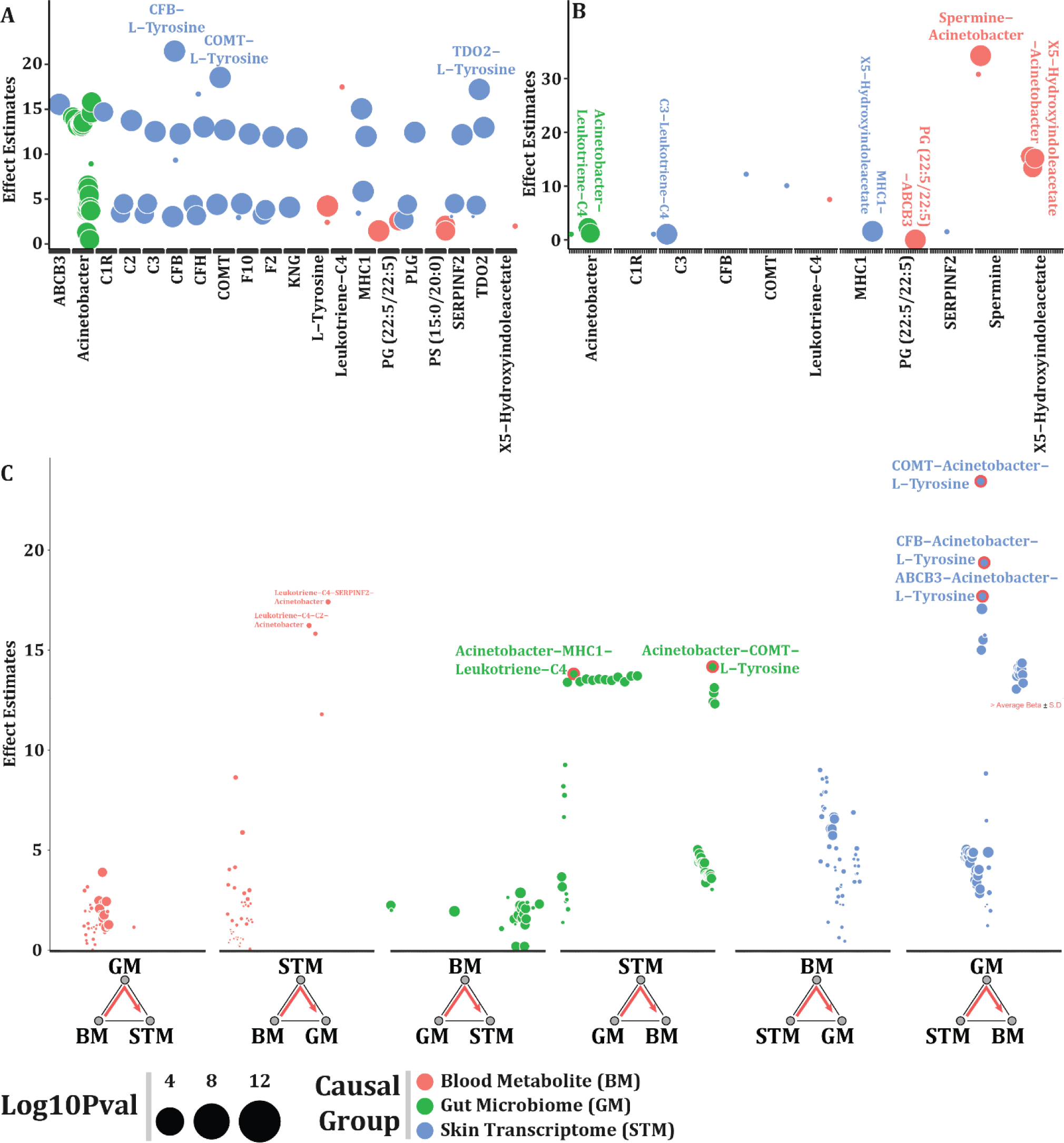
Causal mediation effect on the expression of hub genes underlying variant color morph. A. Average Direct Effect (ADE), illustrating the direct effect of a causal variable on the receptor (e.g., COMT transcriptomic expression on the abundance of Acinetobacter); B. Average Causal Mediation Effect (ACME), illustrating the average effect of mediation variable on the receptor variable (e.g., effect of Spermine on the abundance of Acinetobacter); C. Total effect of causal and mediation effect.

For example, *CFB* and *COMT* expression exhibited the strongest direct causal effect on L-tyrosine concentration via Acinetobacter abundance based on the total effect analysis. Notably, *COMT* showed the highest total effect on L-tyrosine through Acinetobacter. These findings suggest that upregulated *COMT* expression mediated by Acinetobacter abundance might causally influence L-tyrosine concentration. Additionally, both Gut Microbiome -> Skin Transcriptome -> Blood Metabolite and Blood Metabolite -> Skin Transcriptome -> Gut Microbiome linkage direction displayed prominence in the total effect analysis. For instance, Acinetobacter significantly affected L-tyrosine and Leukotriene-C4 concentrations by mediating *COMT* and *MHC1* expression, respectively. Conversely, Leukotriene-C4 had a considerable causal effect on Acinetobacter abundance mediated by *SERPINF2* and *C2* expression.

Interestingly, the average mediation effect (ACME) analysis revealed a different dominant direction: from skin transcriptome to Acinetobacter abundance mediated by blood metabolite concentration (**Fig. 6-B**). Notably, Spermine exhibited the highest ACME on Acinetobacter abundance, suggesting that the interplay between upregulated expression of complement and coagulation cascade genes, particularly *C3*, and Spermine might significantly influence Acinetobacter abundance.

### Interactive Pathways of Gut Microbiome, Blood Metabolome, and Skin Transcriptome

We mapped 329 DMs to the KEGG pathways, among which 206 metabolites were upregulated and 123 metabolites were downregulated in the *TYRP1* mutant compared to the wild WB. We also identified five shared KEGG pathways between DMs and DEGs (**Supplementary Table 1**). Among these pathways, 6 downregulated metabolites, and one upregulated metabolite are involved in the alpha-linolenic acid metabolism, 6 downregulated metabolites in the cysteine and methionine metabolism, 3 downregulated metabolites in the FoxO signaling pathway, 5 downregulated metabolites and 2 upregulated metabolites in the linolenic acid metabolism, and 3 downregulated metabolites in the tyrosine metabolism (**Supplementary Table 1**). We, then, developed a tri-active relationship among gut microbiome, blood metabolome, and transcriptome based on the Pearson correlation matrix. Significant interactions were identified by using a 0.05 FDR cut-off while considering biological reactions according to the Gene-Enzyme-Reaction-Metabolite network based on the MetScape, METACYC, and KEGG database. It has been found that all the mapped pathways are interconnected. Moreover, we identified significant tri-active interactions among gut microbiome, blood metabolites, and skin transcriptomes (**Fig. 7-A-D and Fig. 8-A-B**). We found that the concentration of L-Tyrosine is negatively correlated with the abundance of Acinetobacter and positively correlated with the expression of COMT, which showed a positive correlation with the concentration of L-Normetanephrine (**Fig. 7-B).** According to the Gene-Enzyme-Reaction-Metabolite network based on the MetScape, COMT expression is likely to be overexpressed due to the conversion of L-Tyrosine to L-Normetanephrine via the formation of L-Dopa (**Fig. 8-B**). Moreover, the abundance of Acinetobacter also showed a negative relationship with the concentration of Phosphatidylcholine, which was negatively correlated with the concentration of Arachidonic Acid (**Fig. 7-C**). On the other hand, Arachidonic Acid concentration was positively correlated with the expression of coagulation factor-related genes. However, no significant relations were found in the case of Spermine concentration (**Fig. 7-D**). According to the MetScape, Arachidonic Acid is the end product of the Glycerophospholipid Metabolism and converted to Leukotriene-C4 through Omega-6-Fatty Acid Metabolism (**Fig. 8-B**).

**Fig. 7.**
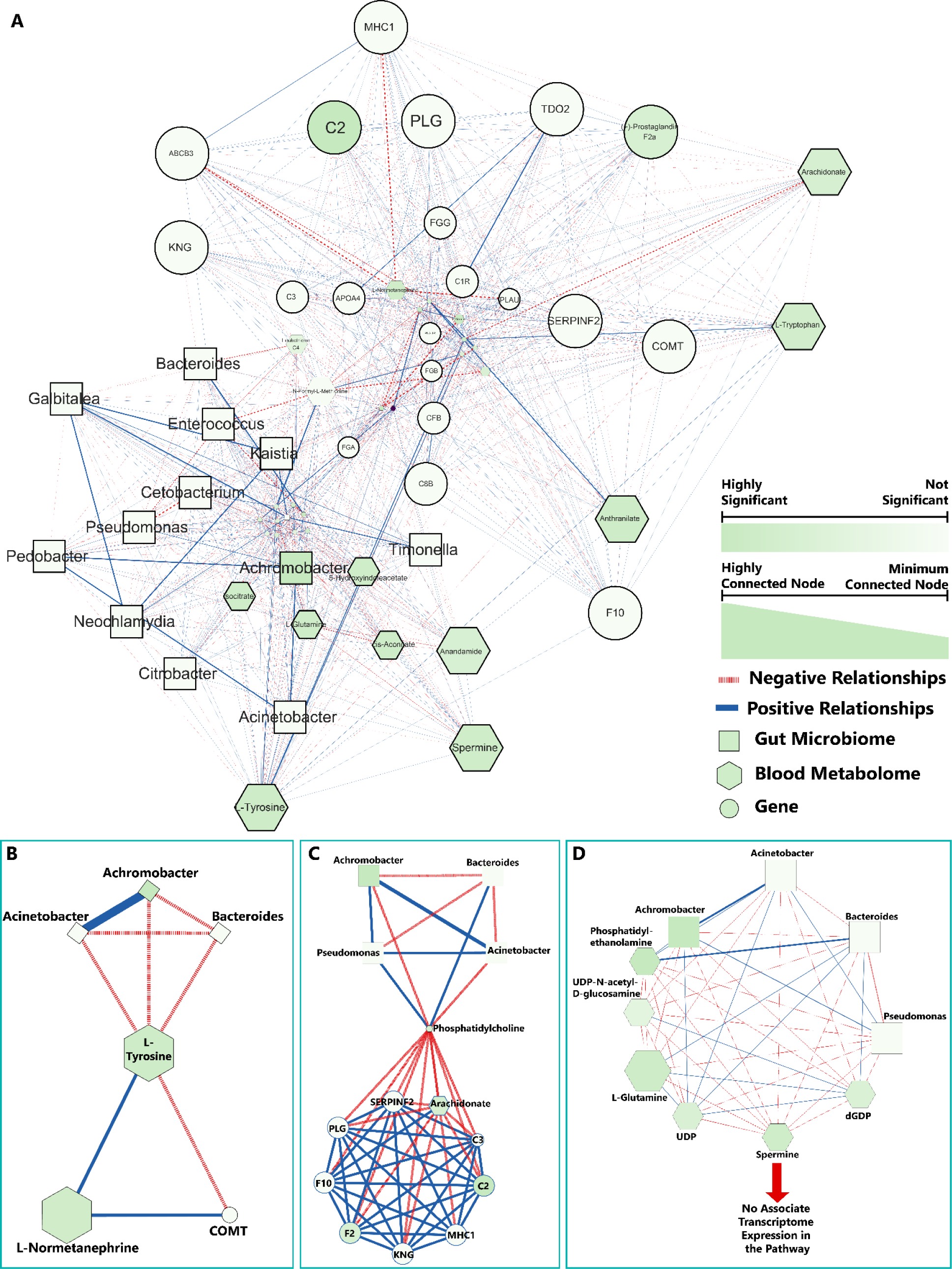
Gut Microbiome-Blood Metabolome-Transcriptome interaction network. A. Tri-active relationship among gut microbiome, blood metabolome, and transcriptome; B. Correlations within the tyrosine metabolism pathway; C. Correlation within the phospholipid metabolism pathway; D. Correlation within polyamine biosynthesis pathway. Microbiome information in the reaction collected from the literature review.

**Fig. 8.**
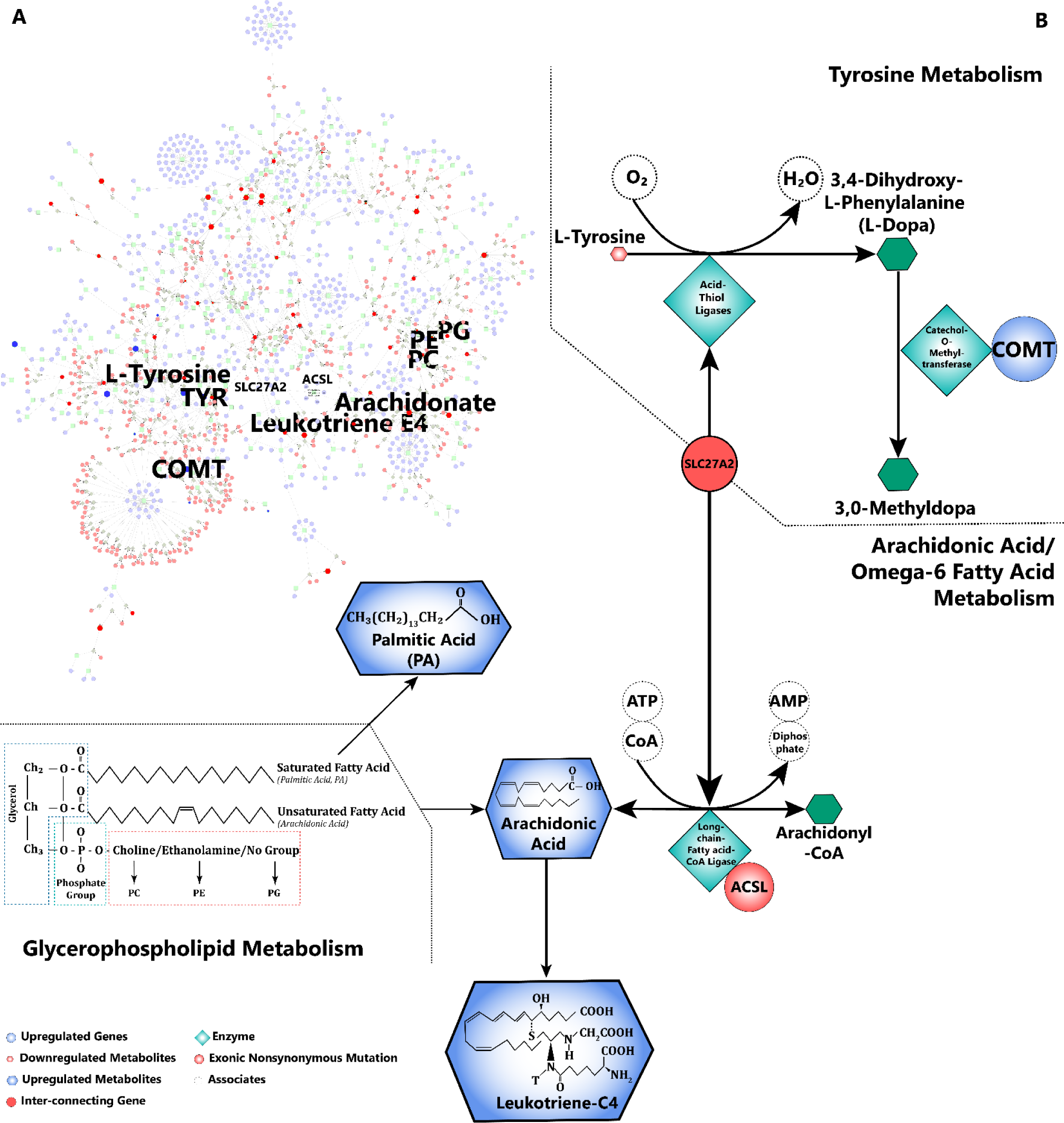
Interactive Pathways of Gut Microbiome, Blood Metabolome, and Skin Transcriptome. A. Pathway-based network of MetScape. B. Gene-Enzyme-Reaction-Metabolite network based on MetScape database, METACYC, and KEGG database

## Discussion

The current study identified that CRISPR/Cas9 mediated *TYRP1*-knocked out genetic variant can significantly regulate an unusual pathogenic gut microbial population, like Acinetobacter. Different discriminating functions, for example, clearly showed that the Acinetobacter population was significantly preferred to be in the *TYRP1* genetic background. We found an elevated expression of *COMT* in the *TYRP1* mutant fishes that have been reported to have a significant causal effect on the abundance of Acinetobacter (U. Boulund et al., 2022; Short, F. L., et al., 2021; Madeo, F. et al., 2018). Gut microbe-associated genes have recently been evident in humans to be involved in immune functions and can influence the abundance of specific microbial communities (U. Boulund et al., 2022). It has also been evident in the in-vitro investigation that *COMT*-derived catecholamines have a significant role in increasing the abundance of pathogenic bacteria, such as Acinetobacter (Belay T. et al., 2002). It is, thus, expected that *TYRP1* may have an anti-microbial role against the abundance of highly pathogenic bacteria and its mutation can regulate this abundance by an upregulated expression of *COMT*.

We observed three metabolic pathways from Acinetobacter-enriched gut to skin of highly expressed anti-inflammatory-related genes (Figure 9). Interaction analysis identified that Acinetobacter shows a significantly positive association with Leukotriene-C4 and Spermine among the top-20 upregulated blood metabolites and negative associations with L-Tyrosine among the top-20 downregulated blood metabolites. Moreover, the total effect mediation analysis determines a significant causal direction from gene expression to blood metabolites mediated by the abundance of Acinetobacter. For example, Acinetobacter has a significant mediation effect on the causal direction from *COMT* expression to the concentration of L-Tyrosine in the *TYRP1* knocked-out genetic background, consistent with previous studies, which suggested that L-Tyrosine can be microbiologically transformed into L-Dopa (Ali S. et al., 2007). It is, thus, expected that *COMT-promoted* Acinetobacter may transform L-Tyrosine to L-Dopa resulting in the highest downregulated blood metabolite. Normetanephrin was also one of the upregulated metabolites in TYRP1 mutant fishes indicating that L-Tyrosine was highly transformed to increase the concentration of Normetanephrin, which has been reported to play an anti-inflammatory role through mediating ABCB3-activated macrophage (Thurm C., et al., 2021; Cristofori F. et al., 2021; Ye, Z. et al., 2020). The total effect mediation analysis also identified that Acinetobacter has a direct causal effect on the concentration of Leukotriene-C4, which has already been identified to cause inflammation (Molina, D. et al., 2007) and derived from the metabolism of Arachidonic acid, which is also one of the top-20 upregulated blood metabolites. Meanwhile, Arachidonic acid is an unsaturated omega-6 fatty acid and is present in phospholipids, including phosphatidylcholine, phosphatidylethanolamine, and phosphatidylinositols, stored in HDL and VLDL (H. Tallima and R. E. Ridi, 2017). It has recently been evident that gut microbial fermentation generates short-chain fatty acids and significantly contributes to hepatic lipid biosynthesis (Yuhan Yin et al., 2022). The current study also identified four significantly upregulated phospholipids (including PG, PC, PS, and PE) that can produce Arachidonic acid catalyzing by phospholipases. It indicates that the upregulated phospholipids carried by HDL and VLDL play a key role in carrying Arachidonic acid and Leukotriene-C4 to the skin, where these metabolites may cause acute inflammation. It is, thus, hypothesized that the COMT-Acinetobacter-Normetanephrin-macrophage driven complement and coagulation cascade-related genes, including *ABCB3, C2, C3, CFB, CFH, F2, F10, KNG, PLG, and TDO2*, were upregulated in the skin in response to an inflammatory effect of Leukotriene-C4 in the *TYRP1* mutant fishes.

**Fig. 9.**
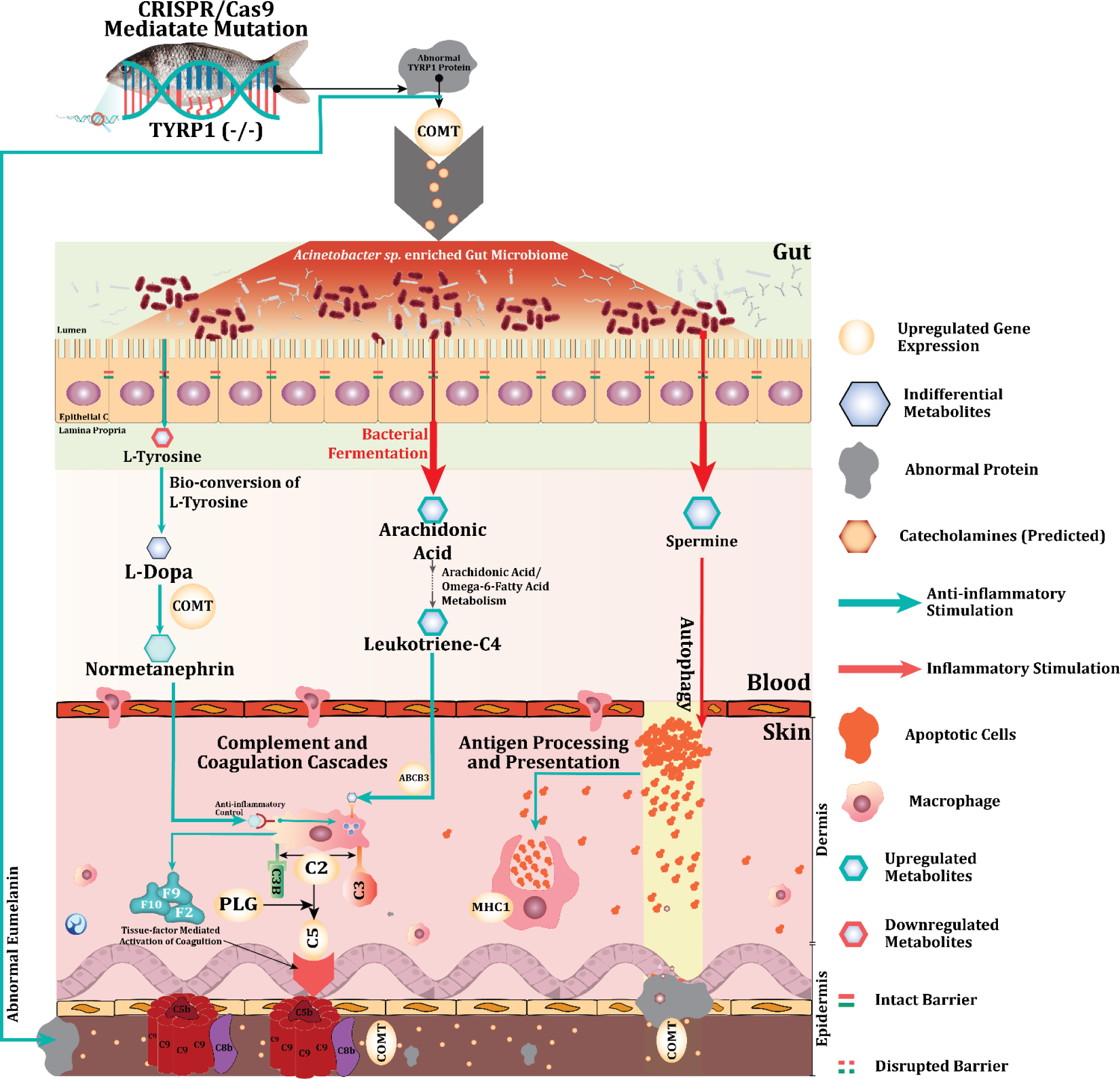
Interplay of gut microbiome, blood metabolome and skin transcriptome under TYRP1 (−/−) genetic background

On the other hand, Spermine is considered one of the natural substrates of Acinetobacter (Short, F. L., et al., 2021; Madeo, F. et al., 2018). In the current study, Spermine has been found to have a significant mediation effect on the abundance of Acinetobacter. This polyamine has also been reported to cause autophagy and break the intact barrier of the epithelial layer of the intestine and even in the skin (Hofer, S.J., et al., 2022; Madeo, F. et al., 2018), which is directly linked with the expression of *C2, C3, CFB, CFH*, and *MHC1* related to complement and coagulation cascades and antigen processing and presentation pathways to engulf the autophagy-derived apoptotic cells. The ACME analysis has shown a similar indication of the causal direction from the upregulated expression of these genes to the abundance of Acinetobacter mediated by the concentration of Spermine in the blood. These findings indicated that a highly upregulated concentration of Spermine may be resulting from Acinetobacter excretion and be a cause of upregulating a set of antigen processing and presentation pathway-related gene expressions to engulf apoptotic cells from Spermine-derived autophagy in the skin.

## Conclusion

The current study intended to make a causal inference among gut microbiome, blood metabolites, and skin transcriptome. We conclude that CRISPR/Cas9 mediated *TYRP1* mutation can derive an auto-immune system that may regulate an unusual pathogenic gut microbiome, Acinetobacter, to play a dual inflammatory role in expressing a set of skin transcriptome underlying unusual coffee-like color phenotype. However, one of the major limitations in this study is the lack of data on gut and skin metabolites, and blood transcriptome. Moreover, the sample size is too small to provide comprehensive inferences of the genetic variants on the causal effect of the gut microbiome on the skin transcriptome mediated by blood metabolites. Further research in vivo with significant samples is needed to disentangle whether the melanin-producing genes have the power to regulate the abundance of gut microbiome leading to transcriptomic expression underlying phenotypic changes in the skin.

## Data availability statement

The raw sequencing reads are available in CNCB (China National Center for Bioinformation) sequence read archives under the Project accession number PRJCA023796 (https://bigd.big.ac.cn/gsa/browse/).

## Funding

The work was supported by the National key research and development program (2022YFD2400102).

## Conflicts of interest

The authors declare that they have no competing interests.

## Author Contributions

Experimental Design: Roland Nathan Mandal, Jun Wang, Chenghui Wang.

Resources, Investigation: Jing Ke, Nusrat Hasan Kanika, Xin Hou

Data Generation: Roland Nathan Mandal, Jing Ke, Zhiyi Zhang, Penghui Zhang, Huifan Chen, Chunxiao Zeng.

Statistical Analysis and Data Interpretation: Roland Nathan Mandal.

Manuscript writing: Roland Nathan Mandal.

Critical Review and Manuscript Edits: Jun Wang, Xiaowen Chen, Chenghui Wang.

